# Woody species with high wood density are more vulnerable to global change in South China

**DOI:** 10.1101/2023.11.06.565905

**Authors:** Guolin C Li, Stavros D Veresoglou

## Abstract

To cope with global change, plants shift their distributions. Distribution shifts tend to be more dramatic across rare species. We here questioned how the distribution range of eight rare woody species is changing and how effectively the plants cope with the shift. We further addressed whether plant traits that could predict those parameters. We carried out Maxent Distribution Modelling on species observation records before 1980 under present climatic conditions and four future (CMIP5) scenarios. To assess how effectively plants cope with migration we assessed species observations after 1980. We finally collated plant trait data on three traits. Most distribution ranges expanded northwards. Temperature driven rather than precipitation driven variables described distribution shifts best. Wood density summarized well the susceptibility of those plants to climate change. There are many woody species in tropical and subtropical areas for which we have very little information available. We identified, subject to our small pool of species, a plant trait, wood density, that could summarize responses to global change that could potentially be used as a tool in conservation ecology to prioritize conservation efforts.

## Introduction

A large fraction of plant biodiversity occurs exclusively in biodiversity hotspots that collectively cover less than 1.5% of the land surface (Myers et al. 2000). Plant species maintaining narrow distributions are particularly vulnerable to climate change driven extinctions (Malcolm et al. 2006; Thomas et al. 2004). As environmental conditions change, plants have to move their distribution (i.e. “migrate”) either polewards or towards higher elevations (Chen et al. 2011; Lenoir et al. 2008). Shifting distributions presents, thereby, a mechanism with which woody plants can alleviate the risk of extinction and it should be trickier for species that maintain narrow distribution ranges to take advantage of it (Tomiolo and Ward 2018). The reason why species with narrower distributions are at disadvantage is that geographic barriers such as sea coasts, lakes and altitudinal topographic gradients get more likely to block migration in species with narrow distributions (Record et al. 2013). It is often difficult to tell which species are coping reasonably with climate change and which ones only occur in areas where they cannot sustain long term growth and will eventually go extinct. In these latter cases we say that the plant species are committing to an extinction dept (Halley et al. 2016; Tilman et al. 1994) which in some cases could get only prevented with human intervention (Kuussaari et al. 2009). There has thus been a consensus that distribution modelling should ideally address the distributions of those species (Lomba et al. 2010; Matern et al. 2007). Modelling the distribution of rare species, however, can be extremely tricky, giving rise to what we know as the “rare species modelling paradox”, that we know little on the distribution modelling of the species that we are interested most (Lomba et al. 2010).

Changes in climate do not occur uniformly throughout earth (e.g. Lobell et al. 2011), meaning that each biodiversity hotspot faces unique challenges in relation to risks of plant extinctions (Malcolm et al. 2006). South-Central China represents a hotspot of plant diversity with 5.5 endemic vascular plant species per 100 km^2^ (compared to a global average of about 0.2 endemic vascular plants per 100 km^2^ of terrestrial habitat). Rapid scientific progress over the last years (He et al. 2010) has facilitated monitoring efforts in the region and has opened up opportunities to map local endemic species distributions. At the same time rapid urbanization in South China has increased considerably the pressure on natural habitats leading to fragmentation of many pristine areas (Seto et al. 2000). This could represent a parameter that renders any assessments of the conservation status of the endemic flora urgent. We combine here historic records on occurrences of eight plant species with more recent observations from two sources of information, the Global Biodiversity Information Facility (GBIF) and the National Herbarium Collection (Chinese Virtual Herbarium) through a distribution modelling exercise to address a series of three questions.

First, we address the relative importance of projected global warming over changes in precipitation frequency and intensity in structuring the distribution ranges of our set of eight plant species (i.e. temperature related vs precipitation related predictors of distribution shifts). Being located within the subtropical climatic zone, habitats in South China, receive considerable amounts of precipitation (Trenberth 2011), which should rarely limit plant growth. At the same time, temperature is changing fast in the region (Stuecker et al. 2020) potentially compromising the ability of some species to persevere in their former distribution range. Unlike herbaceous plants which depend mainly on precipitation, woody species respond strongly to changes in temperature (e.g. Shi et al. 2021; Thurm et al. 2018). This could be why we so often observe woody species to migrate polewards and to higher latitudes (e.g. Lenoir et al. 2008). We thus hypothesize that it will be mainly temperature related variables that drive the distribution of our eight species in our exercise (*Hypothesis One*).

We further question how climate change has altered the distribution range of those eight species and whether the plant species are coping reasonably with this change. All eight plant species in our analysis, describing woody species, share relatively large generation times and got chosen because they maintain narrow distribution ranges. There have been arguments that it is the combination of these two traits that maximize the likelihood that plant species have already committed to an extinction debt (e.g. Kuussaari et al. 2009), which could have been the case for all our eight plant species. Climate change models in the Fifth Assessment Report of IPCC (i.e. Intergovernmental Panel on Climate Change), however, only predict for the subtropical China by the year 2050 moderate increases in temperature of about 1.2°C and increases in precipitation of about 3.4% (Fick and Hijmans 2017). Hence, there is a good chance that any changes in distribution are relatively subtle and that many of these plant species, even those with small distributions, are coping with climate change better than their counterparts close to the poles. We thus, hypothesize, that the location and the size (in sq.km) of their distribution for several of the eight species has not changed much and that the plants have effectively caught up with those changes (*Hypothesis Two*).

We finally collated information on three plant traits for the eight woody species to identify whether there could be some plant traits that predict migration success in plants in this region of the world. Migration success has been linked in the past with several plant traits such as seed mass (Veresoglou and Halley 2018), longevity (Noh et al. 2019; Vellend et al. 2006), pollination strategy and tolerance to external stresses (Saar et al. 2012). At the same time there appears to be a strong positive relationship between tree height and dispersal distances (Thomson et al. 2018), meaning that across natural forests, tree height could potentially predict the success with which trees establish to new habitats. Plant height also peaks at warmer regions (Mao et al. 2019), which could be suggestive of a temperature dependency across woody plants. We thus hypothesized (*Hypothesis Three*) that an easy to collect trait, tree height, effectively captures migration efficiency in our set of eight plant species.

## Materials and Methods

### Plant Species Selection Criteria

We chose eight terrestrial plant species that met the four following criteria:

1. species had been reported in Huang et al. (2017) as occurring exclusively (i.e., being endemic) to two (Geo3 and Geo4) out of the seven geographical regions of China (Liu 1998), covering the center and south of China (15194 out of 18157 species).
2. they described woody species, either trees or shrubs (6489 out of 15194 species).
3. there was a minimum of 20 records on them in the Global Biodiversity Information Facility (GBIF; https://www.gbif.org/).
4. the species had been reported at the Heishiding reserve (23.27° N, 111.53°E) and were thus all describing native late successional forest species (8 out of 6489 species).We integrated this latter criterion to control for the likely inclusion of invasive species or other fast growing species which could have presented idiosyncratic distributions in the area.

The final list comprised eight species. The eight species were the following: *Artocarpus hypargyreus*, *Diospyros strigosa*, *Huodendron biaristatum*, *Machilus breviflora*, *Machilus suaveolens*, *Rhaphiolepis ferruginea*, *Symplocos congesta*, and *Xanthophyllum hainanense*. Out of these species only *Artocarpus hypargyreus* is reported in the International Union for Conservation of Nature (IUCN) list as endangered. We extracted all available records on the eight plant species from GBIF. Four of them had less than 80 complete (i.e. including coordinates and year of the observations) records in GBIF. To increase the number of observations for the subset of the four plants for which we retrieved from GBIF less than 80 records, we further searched the Chinese Virtual Herbarium (CVH; https://www.cvh.ac.cn/) for records that had not been included in GBIF. We retrieved this way between 42 and 192 records per species (i.e. 663 observations in total).

### Environmental variables

We used as predictors the 19 bioclimatic variables that are described in WorldClim (https://worldclim.org/) version 1.4 (Hijmans et al. 2005), presenting averages on climatic variables over the period 1960 – 1990. A complementary non-climatic predictor that can shape the distribution of plant species is altitude (Korner 2007; Lenoir et al. 2008). We thus expanded the set of 19 bioclimatic variables with elevation data (Fick and Hijmans 2017). For all variables we used raster files at a resolution of 30 seconds. To account for collinearities in our observation area which covered south-middle China (18°10’ - 36°22’ N, 97°21’ - 122°43’ E) we quantified correlations between environmental data with the R package “ENMTools” (version 1.0.6; Warren et al. 2021). We set an exclusion threshold of Pearson correlations with coefficients |r| > 0.75 (Dormann et al. 2013; Merow et al. 2013). For correlations of any two climatic variables that were above the threshold, we removed from our dataset the bioclimatic variable with a higher incremental number (i.e. each bioclimatic variable has been assigned a unique ID ranging from 1 to 19). Through this approach we filtered out 12 bioclimatic variables. The bioclimatic variables that remained after this filtering step were the following eight variables: BIO1, BIO2, BIO3, BIO7, BIO12, BIO14, BIO18 and elevation.

### Plant traits

Because of the narrow distributions of the eight species, it proved hard to extract trait information on them. We worked on three traits for which we found data for all eight plant species:

1. *Tree height*: The plant trait summarized the ability of a plant species to intercept light in a close canopy but could also be suggestive of the rooting depth of a plant (Brando 2018). We extracted height values from the Encyclopedia of Life (https://eol.org/).
2. *Leaf size*: we calculated leaf size as the product between width and length of the leaf which we then multiplied with the correcting factor 2/3 (Schrader et al. 2021). Aside presenting an important trait for plant thermoregulation and photosynthetic potential (Leigh 2022), it presents a good predictor of net primary productivity (e.g. Li et al. 2020). We extracted leaf width and length data from the Encyclopedia of Life (https://eol.org/).
3. *Wood density* information at a genus level: The trait shows a high degree of phylogenetic conservatism (Kraft et al. 2010) and thus using data at higher taxonomic levels should pose no major issues. Less than 20% of total variance in wood density occurs within the genus level (Flores and Coomes 2010). We used the median trait values reported at a genus level at ICRAF database from the World Agroforestry Center (http://db.worldagroforestry.org/). Wood density represents a good proxy of mortality rates across tree species (Kraft et al. 2010) but also one of the two attributes to which we can decompose the biomass of woody plants (Phillips et al. 2019). Wood density additionally represents the core trait describing the wood economics spectrum (Chave et al. 2009), being a complementary economics spectrum to that of the leaves (Wright et al. 2006b).

### Species distribution models

To optimize feature classes (i.e., linear, quadratic, hinge etc.) and regularization parameters (Merow et al. 2013) for our models we assessed parsimony (AICc values) of all 60 possible combinations of 15 regularization multipliers (i.e. values 0.25, 0.5, 1, 1.25, 1.5, 1.75, 2, 2.25, 2.5, 2.75, 3, 3.25, 3.5, 3.75, and 4) and 4 feature combinations (i.e. linear, quadratic, hinge, and quadratic with hinge) per species in quintuplicates and extracted mean AICc values of those four runs (Burnham and Anderson 2004; Warren and Seifert 2011). We evaluated the models with ENMeval version 2.0.3 (Muscarella et al. 2014). We used for maximum entropy modelling the model with the lowest AICc value in each set of species per scenario.

We used MaxEnt 3.4.0 (Phillips et al. 2017) to fit the distribution models with 100 bootstrap replicate runs for each species and scenario (Lin et al. 2020; Wang et al. 2022). We used a random subset of 75% of the occurrence data for model training and the other 25% for validation (Garcia et al. 2013; Huberty 1994). To parametrize the near-current prediction algorithms, we used exclusively observations that were taken before the year 1980. To estimate future distribution ranges, we used projections for the bioclimatic variables in the year 2050 based on four global change models in the Climate Model Intercomparison Project 5 (CMIP5): CCSM4, HadGEM2-AO, IPSL-CM5A-LR and MRI-CGCM3 (Yigini and Panagos 2016) under the continue as usual climatic scenario, Representative Concentration Pathways 8.5 (RCP8.5). We extracted the data from WorldClim version 1.4 (https://www.worldclim.org/; Hijmans et al. 2005).We kept the model estimates from the original models depicting current distributions and projected them over each of the future scenarios for the year 2050. We then averaged occurrence probabilities across the four scenarios (Ding et al. 2022).

To assess the quality of our models we used the AUC (Area Under Curve) criterion with the threshold value of 0.7 describing models with acceptable performance (Swets 1988). Predictions on current and future projected distributions were summarized in the form of suitability values per cell ranging between 0 and 1. We classified these values into four ranks: ‘high’ (> 0.6), ‘moderate’ (0.4 - 0.6), ‘low’ (0.2 - 0.4) and ‘unsuitable’ (< 0.2) (e.g. Lin et al. 2020; Yang et al. 2013; Zhang et al. 2018). Based on these thresholds, a suitable distribution described areas with suitability values above 0.2, whereas acceptable distribution with suitability values above 0.4. To infer migration we calculated the kernel (i.e. centroids; *C*_cur_, *C*_fut_) of the distribution ranges (Shi et al. 2021; Skov and Svenning 2004; Thurm et al. 2018). We coupled this to cestimating the arithmetic mean (i.e. centroid) of coordinates for our observations after the year 1981 (*C*_obs_). We assessed the degree to which a species effectively (***effective migration ratio***) migrated to ward its new distribution range as follows:

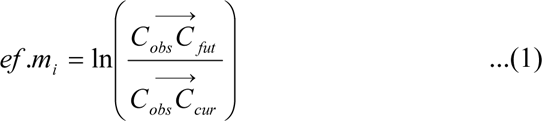

with *i* describing the plant species of interest and C the locations of the respective centroids.

We additionally assessed the ***shift in the distribution range*** as follows:

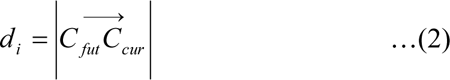

Finally we were interested in the area of the current distributions of species *i*, *S*_i_.

### Statistical analyses

To address whether temperature (rather than precipitation) drives the distribution of our eight species in the specific region of the world (*Hypothesis One*) we quantified the aggregate contribution scores of temperature-related and precipitation-related environmental variables across our eight individual species and compared them with a Mann-Whitney U Test.

To address the degree to which plant species have caught up with any changes of the distribution (*Hypothesis Two*) we calculated the *effective migration ratios* for all species and assessed whether it related (Spearman correlation) to the respective *shifts in their distribution range*. If species had already caught up with the changes in their distribution (*Hypothesis Two*) we should have observed no relationship between these two variables.

To address whether tree height related with migration efficiency (*Hypothesis Three*), we carried out spearman correlations between plant height and the respective *effective migration ratios* as well as with the *shifts in the distribution range*. We also carried out comparable correlation tests with the other plant traits.

In all cases we preferably carried out non-parametric tests. We complemented those with respective parametric tests, even though because of the small pool of plant species in our analyses we could not effectively test the assumption of parametric analyses. All statistical tests were carried on R version 4.2.2 (R core team 2022).

## Results

### Model performance and relationships with environmental variables

Parsimonious model settings (Table S1) across species contained a range of linear, linear quadratic or linear quadratic with hinge features settings and regularization settings varying between 0.25 and 1.75 (we tested 15 regularization settings with values ranging between 0.25 and 4). In all our models the areas under receiver operating characteristic curve (AUCs) exceeded 0.88 (Fig. 1). The lowest AUC value was observed for *Hyodendron biaristatum* (0.881) and the highest for *Diospyros strigosa* (0.981).

**Fig. 1.**
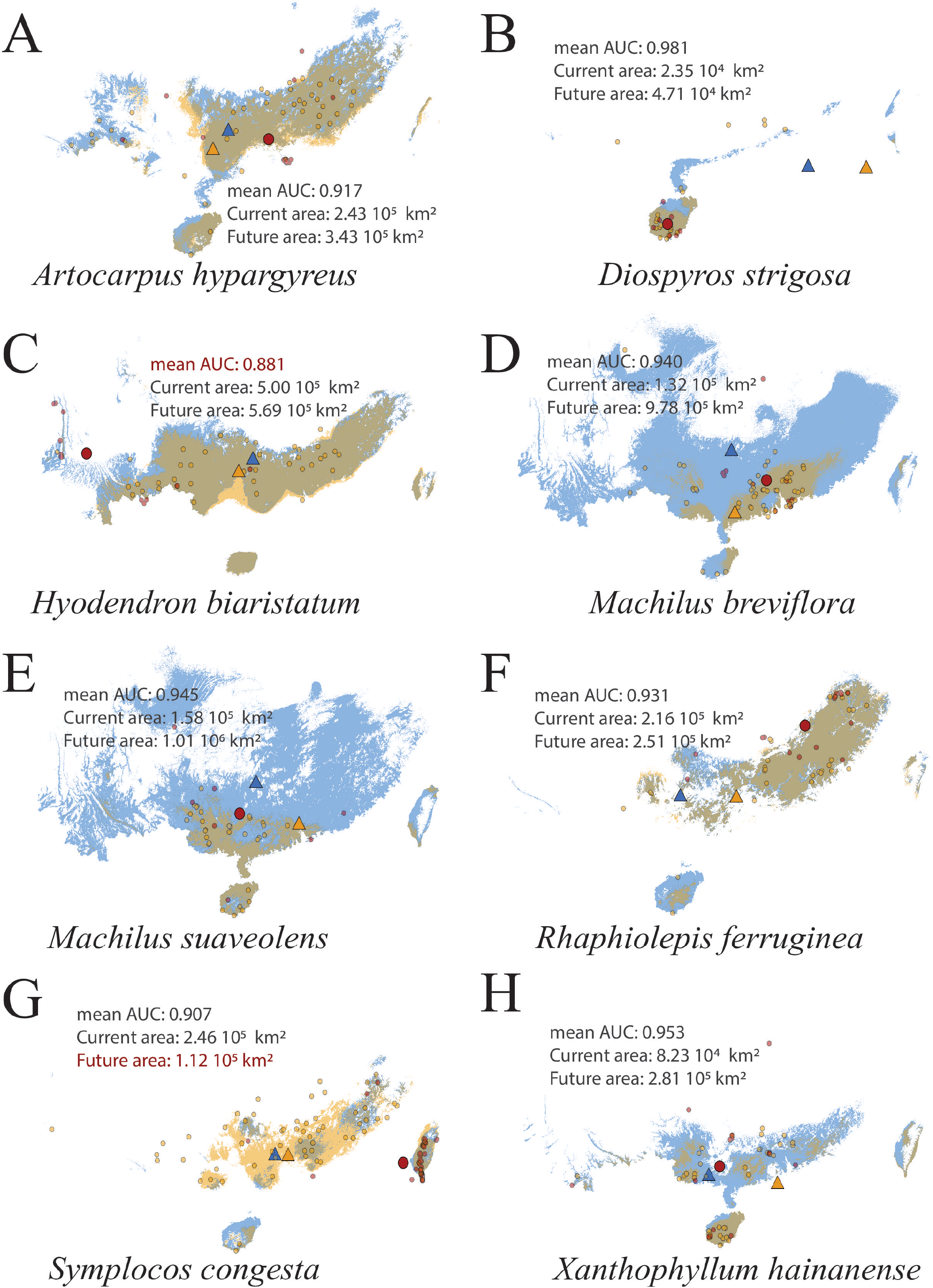
Acceptable (suitability values > 0.4) distributions for the eight species in our study, in relation to present (yellow) and future (blue) climatic conditions. We mark the centroids of the two distributions with large triangles and use a large red circle is to describe the average latitude and longitude of the species observations that were made after 1980. Pre-1980 observations are marked with smaller yellow circles, and post 1980 with red circles. We overlay statistics on model fit and area of the two distributions.

We summarize information on how the environmental variables contributed to the MaxEnt predictions in the form of Table S2. In all eight models the top three most important environmental variables cumulatively accounted for over 73% of the predictive power of the model. In a comparable way the permutation importance proportions of the top three most contributing variables accounted for over 70% across of the total score. The most important environmental variable was annual mean temperature (BIO1; mean contribution 34.66%) and the second most important was annual precipitation (BIO12; mean contribution: 21.54%). Results on the importance of environmental variables were relatively consistent across the different species (Table S2).

The total contribution of the temperature-related bioclimatic variables (BIO1, BIO2, BIO3 and BIO7) for the eight species varied between 35.30% and 93.97% (mean: 69.03% quartiles: 56.16%, 68.70% and 85.92%) The respective contribution estimates for the three precipitation related variables (BIO12, BIO14 and BIO18) were 2.98% - 55.85% and (mean: 27.08%, quartiles: 9.78%, 28.79% and 40.32%). The test between the overall contribution of temperature vs precipitation variables which we carried out to address *Hypothesis One* yielded the following statistics:

### Species geographic distributions

We calculated areas of suitable habitats in different suitability levels of the eight species (Table S3).

*Diospyros strigosa* had the smallest distribution range, whereas *Huodendron biaristatum* the largest (Table S3). 73.8% of the suitable area for *Xanthophyllum hainanense* described occurrence probabilities between 0.2 - 0.4 (i.e. described low suitability area). This proportion was considerably higher than in the other seven species averaging 53.8% of their suitability areas. Our projections to the year 2050 made the forecast that the suitable are of *Xanthophyllum hainanense* will increase by 59.6%. With the exception of *Symplocos congesta*, for which we expect a decline by 2.9% (and a respective decline in the acceptable of 54.4%), we predicted increases in the distribution areas of species for all species. These of the acceptable distributions ranged between 13.9% for *Huodendron biaristatum* and 643.3% for *M. breviflora* (Fig. 1).

We predicted the largest northward shift in the distribution range of a species for *Machilus breviflora* by 3.35 degrees of latitude whereas the respective value for congeneric *Machilus suaveolis* was 2.23 degrees of latitude. On the other extreme, we predicted the smallest northwards swift for *Symplocos congesta* by 0.39 degrees of latitude. We also predicted *Artocarpus hypargyreus* and *Huodendron biaristatum* to move eastwards whereas the other six species westwards. *Xanthophyllum hainanense* evinced the highest westward tendency of centroid movement with 4.19 degrees of longitude (Table S4).

### Wood density predicts migration lag

Post 1980 observations for trees from species for which we had predicted above-average changes in their distributions (historic vs future; here we capture change through the distance to which their distributions changed: variable *shift in the distribution range*) were closer to the historic distribution than in the future one (variable *effective migration ratio*: rho=-0.76, P=0.037; the observation was robust to the consideration of acceptable distributions in which case we also found rho=-0.76, *P*=0.037).

There was no relationship between leaf size or tree height and any of the distribution related variables. We observed, however, that species with a high wood density maintained a smaller distribution area than species with a lighter wood (rho=-0.91, *P*=0.002, Fig. 3a) and were comparatively located closer to their historic distributions than their future ones (relationship with the variable *effective migration ratio*: rho=-0.73, P=0.040, Fig. 3b). The mean wood density across our eight species was 0.657 (quartiles: 0.59, 0.70), above the community weighted value of 0.619 that was reported for tropical systems (Phillips et al. 2019) and above means reported for many common tree families (Fig. 3c). it was species with high wood density that also showed the highest distance change in distribution ranges (relationship with variable *shift in distribution range*: rho=0.90, *P*=0.002).

**Fig. 2.**
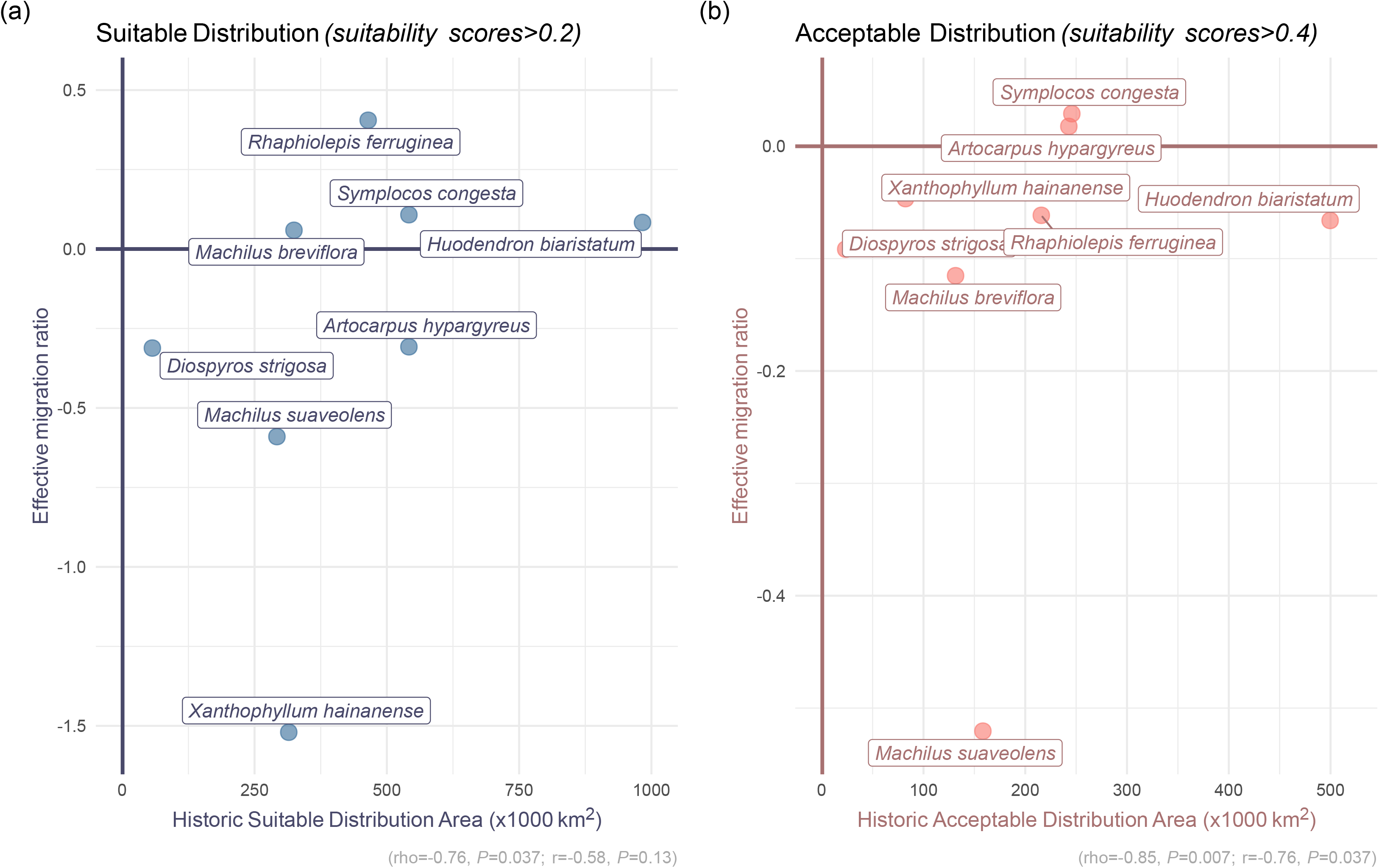
Relationship between the historic distribution area of the eight species and a summary static on the degree to which the most recent observations on a species manifest a high migration efficiency towards the future distribution range (i.e. effective migration ratio – higher values manifest higher efficiency). We summarize the relationships for historic (a) suitable (suitability values > 0.2) and acceptable (suitability values > 0.4) distributions. In both cases we observe positive relationships (but not with parametric statistics in the case of suitable distributions). The take home message is that species that maintained smaller distributions have a harder time catching up with global change.

**Fig. 3.**
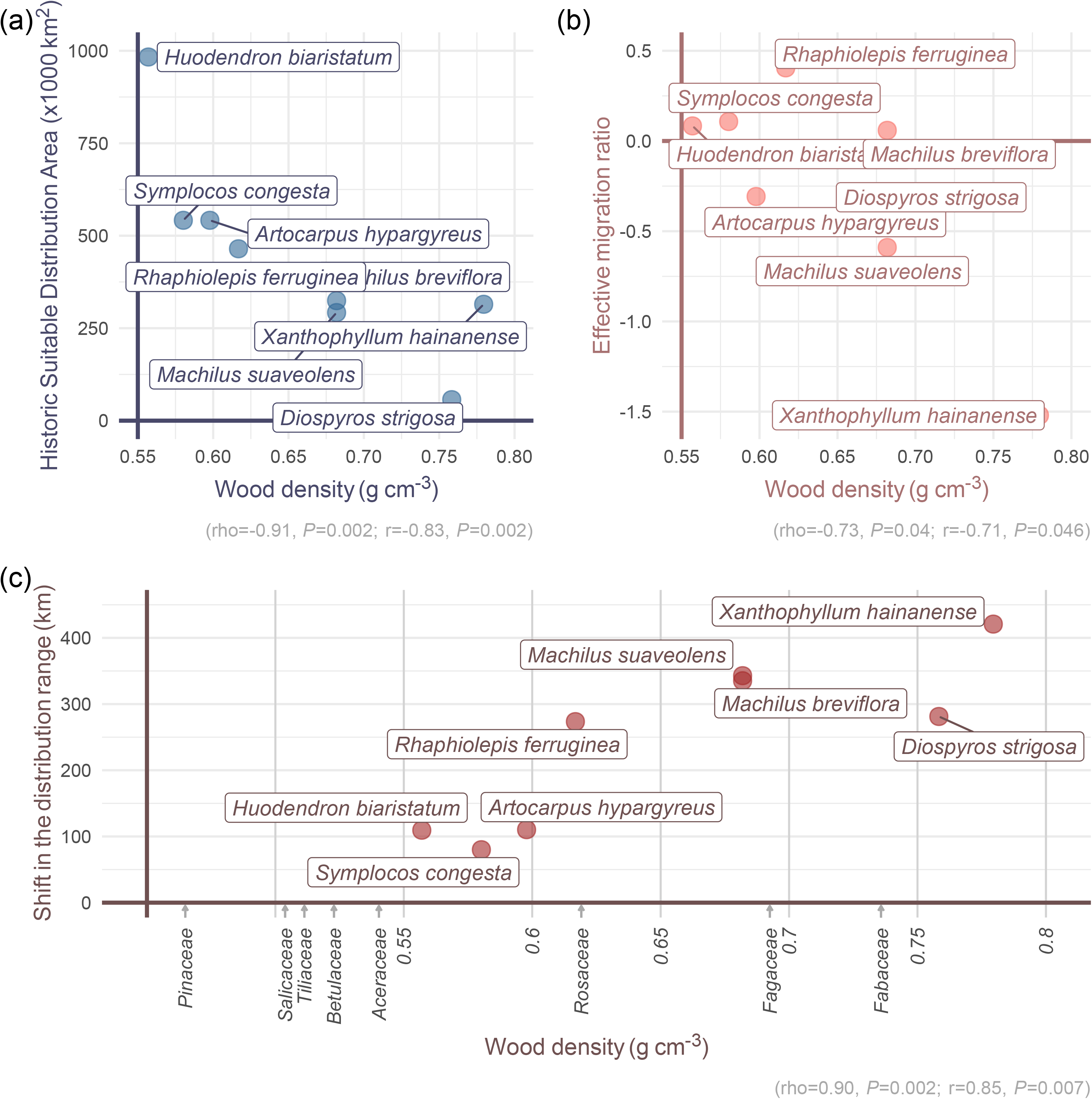
Relationships between wood density (*x*-axis) and (a) historic distribution area; (b) effective migration ratio; (c) the distance between the centroids of future vs historic distributions for the eight plant species. In panel (c) we further present median wood density data for some characteristic families of woody plants implying that we worked with plant species having above average wood density. In all cases we observe strong relationships suggesting that wood density is a plant trait that can effectively predict migration efficiency across woody plant species.

## Discussion

We modeled here the distribution ranges of eight plant species in South China under past and future climatic scenarios to assess how they may respond to anticipated global change. We hypothesized that (*Hypothesis 1*) the distribution ranges are mainly influenced by temperature related rather than precipitation related changes which was the case (Table S2). We further expected (*Hypothesis 2*) that the distribution range of most of the species would be resilient to climate change. We observed universal northward shifts in agreement with many other empirical studies (Kumar and Rawat 2022; Lu et al. 2021; Mueller et al. 2022; Yang et al. 2022) as well as that the distribution ranges of most species increased in size (Fig. 1). We finally hypothesized (*Hypothesis 3*) that we would be able to identify plant traits capturing migration efficiency and observed that species with a higher wood density maintained a smaller distribution area, a higher shift in distribution range and heavier difficulty to cope with climate change among the eight (Fig. 3).

Many of the animals that have gone extinct presented distinct features – traits. As an example dodo had evolved in an environment without natural predators which made it an easy target for human hunters (Roberts and Solow 2003) whereas the thunder bird (*Genyornis newtoni*), another flightless bird in Australia, evolved an island gigantism (Miller et al. 2016). Aside small body size, animals that effectively disperse to larger distances may also be more resilient to climate change (Nadeau and Fuller 2016). It is unclear if comparable traits exist for plants that can summarize their susceptibility to global change. There have been some studies observing correlations between life history traits such as flowing phenology, seed size, stress tolerance, dispersal mode and plant height with extinction risk in plants (Fréville et al. 2007), but these have not been consistent across ecosystems (e.g. Matteodo et al. 2013). O’Rourke et al. (2022) assessed the degree to which the extinction risk state of the Irish Flora could be predicted by a range of fourteen leaf traits to arrive at the conclusion that any such correlation should be weak in magnitude. It might be easier, on the other hand to predict extinction risk, which we earlier claim to relate to migration efficiency for woody plants, instead as a number of studies has already observed correlation with traits linked to migration (Liao et al. 2020), longevity (Noh et al. 2019; Vellend et al. 2006), pollination strategy and tolerance to external stress caused by changes (Saar et al. 2012).

In this study we considered three plant traits for which we could collect data for all eight plant species, leaf size, tree heigh and wood density. We only observed correlations of the migration variables with wood density. The wood economics spectrum has recently proposed as an extension of the leaf economics spectrum (Chave et al. 2009). Trees with a low wood density, for example, may maintain higher relative growth rates but experience a higher mortality (Chave et al. 2009). The likely relationships with growth and mortality rates may be the exact reason why wood density could in our study predict migration parameters. The wood economics spectrum may not be completely independent from the respective leaf’s spectrum (Reich et al. 2003; Zhao et al. 2017), meaning that high wood and leaf densities could compromize growth rates but secure tissue longevity (Wright et al. 2006a). Our focus on woody species may explain why a wood economics spectrum variabe, wood density outperformed variables from the leaf economics spectrum such as leaf size in predicting migration.

Although each of the biodiversity hotspots faces unique challenges in relation to risks of plant extinctions, such as degree of endemism and biome specificity across plant species (Malcolm et al. 2006), our results might be generlizable across natural systems and biodiversity hotspots. First, because we specifically targetted rare, likely endemic to this biodiversity hotspot, plant species which thus experience a high degree of genetic isolation. We believe that this should be descriptive for most rare plant species across all biodiversity hotspots. Second, we assessed migration parameters in relation to three crude plant traits, which capture three common life history syndromes (i.e. wood -wood density- and leaves -leaf size-economic spectrum but also body mass – tree height, for the three traits, respectively) across plant species and should be generalizable across habitats. Plant rarity increases consisderably susceptibility to environmental variability (Malcolm et al. 2006; Thomas et al. 2004). This implies that through our analysis we specifical targeted species that may experience large changes in their distributions in light of global change.

A limitation of this study is that we only worked on the distribution of eight plant species. We rationalize on this on the grounds that these were the only eight woody species that met our filtering criteria. Could, thereby, the relationships that we observed been due to idiosyncrases of thespecies we considered? We observed clear relationships across our variables and paired our analyses with conservative and robust to likely outliers non-parameteric spearman rho tests. Moreover, many of the relationships we observed were in agreement with our expectations. For example, species with a larger distribution have been migrating more effectively (Fig. 2), which is in agreement with the views from Malcolm et al. (2006) that such species are less vulnerable to climate change. It may, neverthess, be difficult to generalize the findings for other regions. As an example, most European wood plant species maintain considerably larger distribution areas. We are, nevertheless, confident that we captured a major driver of susceptibility of woody plants to climate change in the specific region.

Notwithstanding the likely generality of the findings there have been additional reasons why we believe the focus on this specific region was important. We addressed a region where global change models predict strong alterations in precipitation frequency and intensity (e.g. Trenberth 2011) and through our analysis we questioned whether these would overwhelm the anticipated importance of temperature related parameters in relation to susceptibility risk. Moreover, a lot of plants in the region remain undocumented and as many as 794 new plant species are being discovered per year (http://sp2000.org.cn/). This implies that assessing extinction risk in the region for those less well-documented plant species might only be possible through general predictive models which integrate plant traits (McGill et al. 2006).

We present here a distribution range modeling exercise supporting that high wood density across rare plant species contributes to relatively smaller geographic distribution ranges and more pronounced range shifts. Wood density could hence present an inconspicuous constituent of plant physiology linking plant growth components to shifts in distribution ranges and susceptivity to environmental change and could thus open up opportunities for larger syntheses in plant biogeography. One remaining open question is the degree to which our observations might be generalizable across habitats and biomes. At the same time, it remains unclear if through collating information on additional traits we could come up with superior predictors of migration potential across plant species. Irrespective of those two perspectives, replicating the analysis across biodiversity hotspots could contribute immensely in synthesizing across trait and extinction ecology.

## Supplementary materials file

### Appendix 1: Detailed Materials and Methods

We gathered from GBIF and the Chinese Virtual Herbarium (CVH) a total of 420 observations (Table S5). We classified them into the classes of historic observations (prior to 1980) which we modelled with historic climatic settings and recent observations (post 1980) which we used to assess the efficiency with which the plant species catch up with climate change.

To fit our models we used the Java version (MaxEnt.jar) of Maxent v3.4.2. Our models were as follows:

#### 1. Predictors

To address over-fitting, in our models, we excluded collinear environmental variables (Dormann et al. 2013; Merow et al. 2013). Dormann et al. 2013 suggested an absolute threshold of Pearson correlation coefficient 0.70 to exterminate collinearity in most situations. Variance inflation factor (VIF) test is alterative (Naimi et al. 2014). However, if a study has aims, for example, on which variables drive species distribution, Merow et al. 2013 suggested not to prescreen the predictors too excessively. In this study, we set an exclusion threshold of Pearson correlation coefficient |r| > 0.75 to necessarily and not excessively prescreen predictors. Attached to this step, we used the variance inflation factor (VIF) to check collinearity (VIF < 5 for no collinearity, and VIF >10 for significant collinearity), and using this threshold the VIF was still 52. We faced a trade-off on prescreening and remaining variables. Since thresholds from 0.7 to 0.8 were largely used in this field, we considered this level of collinearity acceptable, because we would select relatively best settings to avoid overfitting afterwards.

#### 2. Size of the modelling area

To increase the accuracy of the predictions we restricted the size of the modelling area to tropical, subtropical and a few temperate regions in China. The exact window was as follows: 18°10’ - 36°22’ N, 97°21’ - 122°43’ E.

#### 3. Test set

We consistently, across our models, used a regular set of 75% of observations for training and 25% of observations for validation by randomly sampling in agreement with recommendations in the literature (Garcia et al. 2013; Phillips 2008).

#### 4. Settings of the two parameters

We experimented with the following feature settings: appropriate feature classes (i.e. linear, quadrat and hinge), regularization parameters and (we used 15 different parameters ranging from 0.25 to 4) and nonspatial partitioning techniques (optimal nonspatial partitioning techniques were decided based on the number of observations) (Merow et al. 2013). We decided on optimal models (Table S1) based on parsimony – AICc values.

#### 5. Representative of plant species pool (maxent modelling)

Our pool of wight species presented all eight woody species that met the four species inclusion criteria (main manuscript). We understand that our finding on wood density may be difficult to get generalized across other ecoregions, globally. As an example, most European wood plant species maintain considerably larger distribution areas. We are, nevertheless, confident that we captured a major driver of susceptibility of woody plants to climate change in the specific region.

---

**Table S1.**
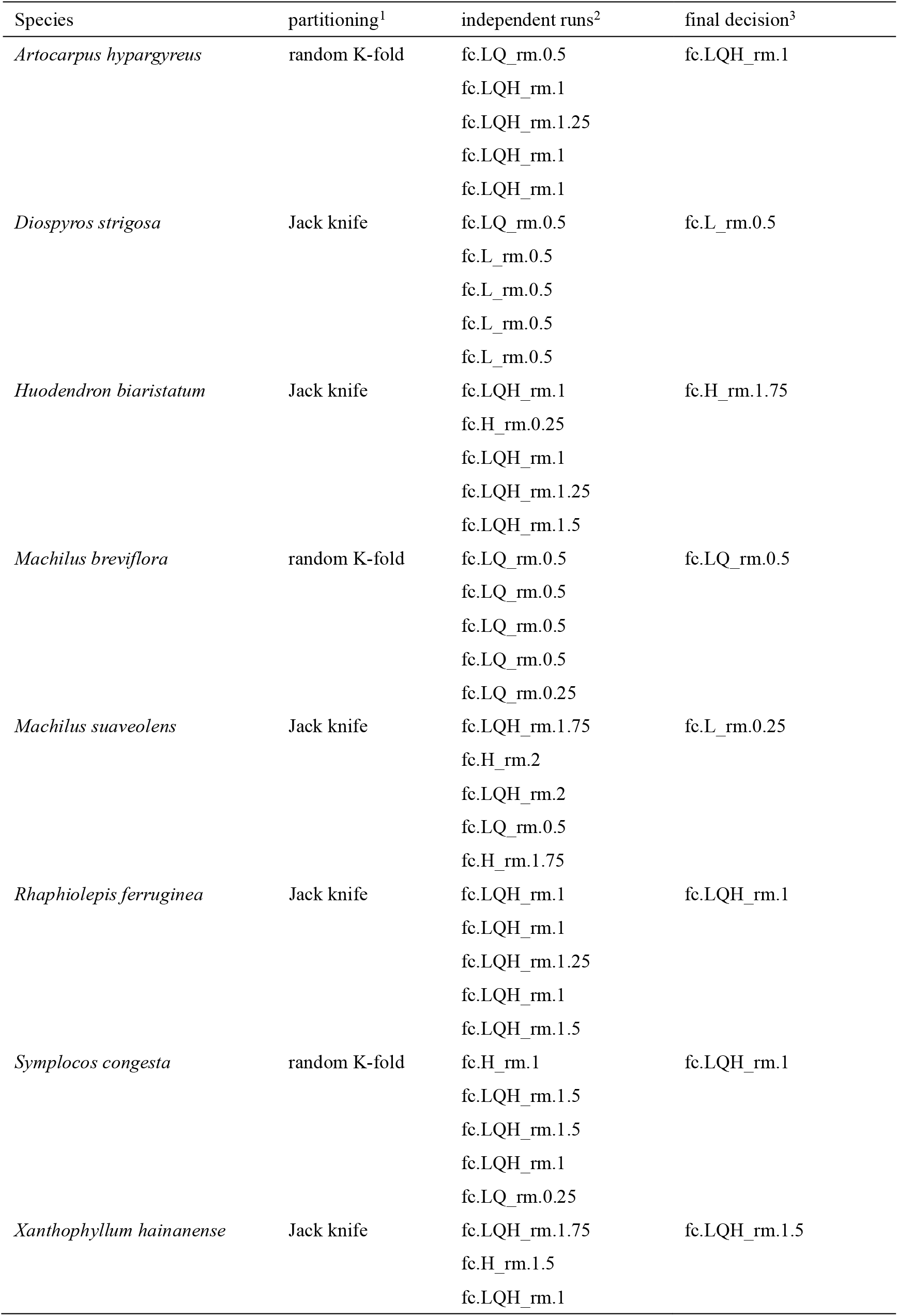
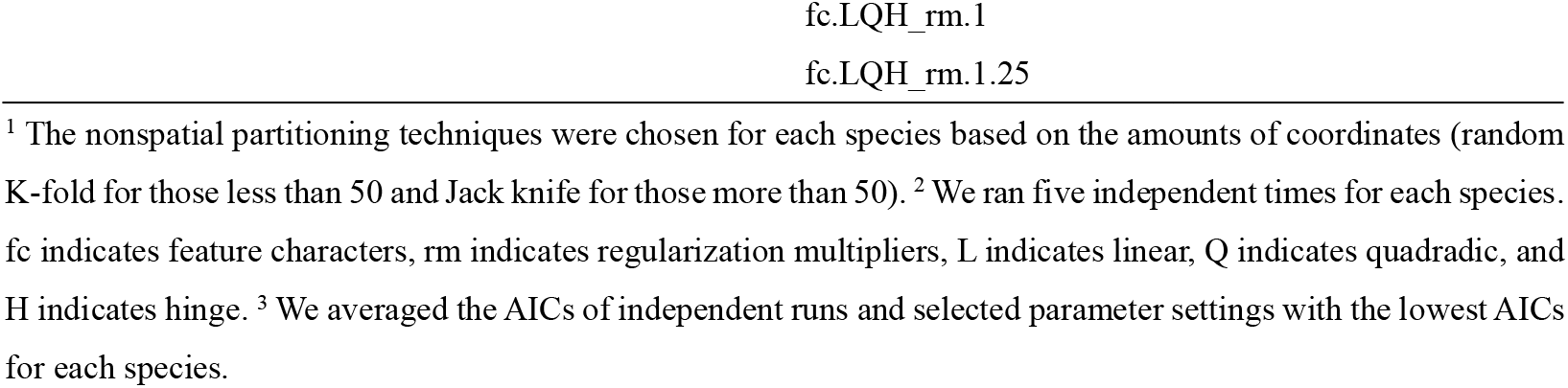
Parsimonious model settings across species contained a range of linear, quadratic, hinge or quadratic with hinge features settings and regularization settings varying between 0.25 and 1.75 (which was decided after testing 15 regularization settings ranging between 0.25 and 4) were selected based on non-duplicate coordinate data.

| Species | partitioning<sup>1</sup> | independent runs<sup>2</sup> | final decision<sup>3</sup> |
| --- | --- | --- | --- |
| *Artocarpus hypargyreus* | random K-fold | fc.LQ_rm.0.5<br>fc.LQH_rm.1<br>fc.LQH_rm.1.25<br>fc.LQH_rm.1<br>fc.LQH_rm.1 | fc.LQH_rm.1 |
| *Diospyros strigosa* | Jack knife | fc.LQ_rm.0.5<br>fc.L_rm.0.5<br>fc.L_rm.0.5<br>fc.L_rm.0.5<br>fc.L_rm.0.5 | fc.L_rm.0.5 |
| *Huodendron biaristatum* | Jack knife | fc.LQH_rm.1<br>fc.H_rm.0.25<br>fc.LQH_rm.1<br>fc.LQH_rm.1.25<br>fc.LQH_rm.1.5 | fc.H_rm.1.75 |
| *Machilus breviflora* | random K-fold | fc.LQ_rm.0.5<br>fc.LQ_rm.0.5<br>fc.LQ_rm.0.5<br>fc.LQ_rm.0.5<br>fc.LQ_rm.0.25 | fc.LQ_rm.0.5 |
| *Machilus suaveolens* | Jack knife | fc.LQH_rm.1.75<br>fc.H_rm.2<br>fc.LQH_rm.2<br>fc.LQ_rm.0.5<br>fc.H_rm.1.75 | fc.L_rm.0.25 |
| *Raphiolepis ferruginea* | Jack knife | fc.LQH_rm.1<br>fc.LQH_rm.1<br>fc.LQH_rm.1.25<br>fc.LQH_rm.1<br>fc.LQH_rm.1.5 | fc.LQH_rm.1 |
| *Symplocos congesta* | random K-fold | fc.H_rm.1<br>fc.LQH_rm.1.5<br>fc.LQH_rm.1.5<br>fc.LQH_rm.1<br>fc.LQ_rm.0.25 | fc.LQH_rm.1 |
| *Xanthophyllum hainanense* | Jack knife | fc.LQH_rm.1.75<br>fc.H_rm.1.5<br>fc.LQH_rm.1 | fc.LQH_rm.1.5 |

<sup>1</sup> The nonspatial partitioning techniques were chosen for each species based on the amounts of coordinates (random K-fold for those less than 50 and Jack knife for those more than 50). <sup>2</sup> We ran five independent times for each species. fc indicates feature characters, rm indicates regularization multipliers, L indicates linear, Q indicates quadratic, and H indicates hinge. <sup>3</sup> We averaged the AICs of independent runs and selected parameter settings with the lowest AICs for each species.

**Table S2.** Contribution of environmental variables^1^ to optimal models for the eight species.

| Species |  | BIO1 | BIO2 | BIO3 | BIO7 | BIO12 | BIO14 | BIO18 | elev |
| --- | --- | --- | --- | --- | --- | --- | --- | --- | --- |
| *Artocarpus hypargyreus*<br>tree | C<sup>2</sup> | 27.87 |  | 28.16 |  | 28.99 |  |  |  |
|  | PI<sup>3</sup> | 39.68 |  | 15.20 |  | 28.42 |  |  |  |
|  | JR<sup>4</sup> | I |  | II |  | III |  |  |  |
| *Diospyros strigosa*<br>shrub | C | 28.37 |  |  | 64.92 |  |  | 3.12 |  |
|  | PI | 36.41 |  |  | 23.72 | 15.87 |  |  |  |
|  | JR | I | III |  | II |  |  |  |  |
| *Huodendron biaristatum*<br>shrub | C | 17.54 |  | 31.82 |  | 30.73 |  |  |  |
|  | PI |  | 40.72 | 17.08 | 16.32 |  |  |  |  |
|  | JR | I |  | II |  | III |  |  |  |
| *Machilus breviflora*<br>tree | C | 56.73 |  | 7.37 |  | 17.30 |  |  |  |
|  | PI | 43.62 |  | 11.77 |  | 20.98 |  |  |  |
|  | JR | I |  |  |  | II |  | III |  |
| *Machilus suaveolens*<br>tree | C | 75.33 |  |  | 18.23 |  |  |  | 1.62 |
|  | PI | 72.08 | 6.24 |  |  |  |  |  | 14.38 |
|  | JR | I |  |  | II |  |  |  | III |
| *Raphiolepis ferruginea*<br>shrub | C | 10.91 |  | 20.71 |  | 41.99 |  |  |  |
|  | PI |  | 23.09 |  |  | 29.67 |  |  | 10.62 |
|  | JR | III |  | II |  | I |  |  |  |
| *Symplocos congesta*<br>tree | C |  |  | 15.84 | 22.39 | 38.94 |  |  |  |
|  | PI |  |  | 12.99 | 25.03 | 32.34 |  |  |  |
|  | JR | II |  |  | III | I |  |  |  |
| *Xanthophyllum hainanense*<br>tree | C | 49.73 |  |  | 15.32 | 12.41 |  |  |  |
|  | PI | 37.28 |  |  |  | 14.68 |  |  | 17.47 |
|  | JR | I |  | III | II |  |  |  |  |

<sup>1</sup> The top three contributing variables for each species were displayed. <sup>2</sup> C: contribution (%). <sup>3</sup> PI: permutation importance (%). <sup>4</sup> JR: Jackknife of regularized training gain, and roman numerals indicate the contribution ranks.

**Table S3.** Areas of the suitable habitats in different suitability levels.

| Areas (10000 km<sup>2</sup>) | Year | low | moderate | high | acceptable | suitable |
| --- | --- | --- | --- | --- | --- | --- |
| *Artocarpus hypargyreus* | 1980 | 29.88597 | 18.21396 | 6.068194 | 24.28215 | 54.16812 |
| *Diospyros strigosa* | 1980 | 3.330417 | 1.432431 | 0.923125 | 2.355556 | 5.685972 |
| *Huodendron biaristatum* | 1980 | 48.35299 | 31.97 | 17.99056 | 49.96056 | 98.31354 |
| *Machilus breviflora* | 1980 | 19.29625 | 8.495278 | 4.6575 | 13.15278 | 32.44903 |
| *Machilus suaveolens* | 1980 | 13.38785 | 10.20396 | 5.632708 | 15.83667 | 29.22451 |
| *Raphiolepis ferruginea* | 1980 | 24.89111 | 16.87493 | 4.702292 | 21.57722 | 46.46833 |
| *Symplocos congesta* | 1980 | 29.59187 | 21.14285 | 3.434097 | 24.57694 | 54.16882 |
| *Xanthophyllum hainanense* | 1980 | 23.23222 | 6.135903 | 2.093542 | 8.229444 | 31.46167 |
| *Artocarpus hypargyreus* | 2050 | 42.31951 | 27.56458 | 6.705625 | 34.27021 | 76.58972 |
| *Diospyros strigosa* | 2050 | 10.65146 | 3.212222 | 1.496319 | 4.708542 | 15.36 |
| *Huodendron biaristatum* | 2050 | 67.83431 | 51.97979 | 4.946528 | 56.92632 | 124.7606 |
| *Machilus breviflora* | 2050 | 42.275 | 33.88174 | 63.88701 | 97.76875 | 140.0437 |
| *Machilus suaveolens* | 2050 | 46.87972 | 35.27931 | 66.02396 | 101.3033 | 148.183 |
| *Raphiolepis ferruginea* | 2050 | 24.97562 | 19.15986 | 5.983958 | 25.14382 | 50.11944 |
| *Symplocos congesta* | 2050 | 41.42312 | 8.859792 | 2.338125 | 11.19792 | 52.62104 |
| *Xanthophyllum hainanense* | 2050 | 22.08646 | 17.76285 | 10.36667 | 28.12951 | 50.21597 |

**Table S4.** Geographic coordinates of the distribution centroids in near-current predictions and future projections.

| Species | 1980 predictions |  | after 1980 |  | 2050 projections |  |
| --- | --- | --- | --- | --- | --- | --- |
|  | longitudes | latitudes | longitudes | latitudes | longitudes | latitudes |
| *Artocarpus hypargyreus* | 110.57 | 22.859 | 113.2948 | 23.23373 | 111.3198 | 23.66754 |
| *Diospyros strigosa* | 119.2115 | 21.38312 | 109.5565 | 18.88741 | 116.3994 | 21.43851 |
| *Huodendron biaristatum* | 109.5419 | 24.10164 | 100.2931 | 25.01755 | 110.3949 | 24.78714 |
| *Machilus breviflora* | 110.768 | 22.00353 | 112.7116 | 23.67576 | 110.5679 | 25.3497 |
| *Machilus suaveolens* | 113.4465 | 23.26277 | 109.8554 | 23.73984 | 110.8393 | 25.49018 |
| *Raphiolepis ferruginea* | 112.607 | 23.30669 | 115.9551 | 26.33625 | 109.8727 | 23.36253 |
| *Symplocos congesta* | 112.6681 | 23.45628 | 119.6794 | 22.99352 | 111.8691 | 23.49478 |
| *Xanthophyllum hainanense* | 113.0054 | 21.66383 | 109.491 | 22.51431 | 108.8197 | 22.09638 |

**Table S5.** Analytical statistics on the observations we gathered per species.

| Species | GBIF | CVH | all_obs | before1980 |  | after1981 |  |
| --- | --- | --- | --- | --- | --- | --- | --- |
|  |  |  |  | collected | used | collected | used |
| *Artocarpus hypargyreus* Hance | 85 |  | 85 | 67 | 56 | 18 | 17 |
| *Diospyros strigosa* Hemsl. | 25 | 17 | 42 | 35 | 27 | 7 | 7 |
| *Huodendron biaristatum* Rehder | 82 |  | 82 | 61 | 42 | 21 | 21 |
| *Machilus breviflora* (Benth.) Hemsl. | 52 | 38 | 90 | 74 | 67 | 16 | 12 |
| *Machilus suaveolens* S.K.Lee | 22 | 24 | 46 | 38 | 36 | 8 | 7 |
| *Raphiolepis ferruginea* Metcalf | 32 | 18 | 50 | 34 | 30 | 16 | 16 |
| *Symplocos congesta* Benth. | 192 |  | 192 | 34 | 72 | 98 | 95 |
| *Xanthophyllum hainanense* Hu | 36 | 40 | 76 | 59 | 43 | 17 | 14 |

**Appendix II:** Raw data.

| species | acceptableLogRR | fullLogRR | EquilateRatioFull | DistributionChangeFull | EquilateRatioAcceptable | DistributionChangeAcceptable | FullDistributionArea | AcceptableDistributionArea | n | woodDensity | height | LeafSize |
| --- | --- | --- | --- | --- | --- | --- | --- | --- | --- | --- | --- | --- |
| *Artocarpus hypargyreus* | 0.34 | 0.35 | -0.31 | 1.10 | 0.02 | 1.00 | 54.17 | 24.28 | 85 | 0.60 | 20 | 55.00 |
| *Diospyros strigosa* | 0.69 | 0.99 | -0.31 | 2.81 | -0.09 | 1.49 | 5.69 | 2.36 | 42 | 0.76 | 8 | 37.33 |
| *Huodendron biaristatum* | 0.13 | 0.24 | 0.08 | 1.09 | -0.07 | 0.78 | 98.31 | 49.96 | 82 | 0.56 | 22 | 48.17 |
| *Machilus breviflora* | 2.01 | 1.46 | 0.06 | 3.35 | -0.12 | 2.34 | 32.45 | 13.15 | 90 | 0.68 | 8 | 5.83 |
| *Machilus suaveolens* | 1.86 | 1.62 | -0.59 | 3.43 | -0.52 | 3.36 | 29.22 | 15.84 | 46 | 0.68 | 7 | 27.73 |
| *Raphiolepis ferruginea* | 0.15 | 0.08 | 0.41 | 2.73 | -0.06 | 0.19 | 46.47 | 21.58 | 50 | 0.62 | 10 | 40.00 |
| *Symplocos congesta* | -0.79 | -0.03 | 0.11 | 0.80 | 0.03 | 0.26 | 54.17 | 24.58 | 192 | 0.58 | 2 | 26.67 |
| *Xanthophyllum hainanense* | 1.23 | 0.47 | -1.52 | 4.21 | -0.05 | 0.41 | 31.46 | 8.23 | 76 | 0.78 | 20 | 26.00 |

## References

Brando P (2018) Tree height matters. Nat Geosci 11:390–391

Burnham KP, Anderson DR (2004) Multimodel inference - understanding AIC and BIC in model selection. Sociol Methods Res 33:261–304

Chave J, Coomes D, Jansen S, Lewis SL, Swenson NG, Zanne AE (2009) Towards a worldwide wood economics spectrum. Ecol Lett 12:351–366

Chen IC, Hill JK, Ohlemuller R, Roy DB, Thomas CD (2011) Rapid Range Shifts of Species Associated with High Levels of Climate Warming. Science 333:1024–1026

Ding WC, Li HY, Wen JB (2022) Response of the invasive plant Ailanthus altissima (Mill.) Swingle and its two important natural enemies (Eucryptorrhynchus scrobiculatus (Motschulsky) and E. brandti (Harold)) to climate change. Ecol Indic 143:14

Dormann CF, Elith J, Bacher S, Buchmann C, Carl G, Carre G, Marquez JRG, Gruber B, Lafourcade B, Leitao PJ, Munkemuller T, McClean C, Osborne PE, Reineking B, Schroder B, Skidmore AK, Zurell D, Lautenbach S (2013) Collinearity: a review of methods to deal with it and a simulation study evaluating their performance. Ecography 36:27–46

Fick SE, Hijmans RJ (2017) WorldClim 2: new 1-km spatial resolution climate surfaces for global land areas. Int J Climatol 37:4302–4315

Flores O, Coomes DA. 2010. Estimating the wood density of species for carbon stock assessments. Methods in Ecology and Evolution 2: 214–220.

Fréville H, McConway K, Dodd M, Silvertown J (2007) Prediction of extinction in plants: Interaction of extrinsic threats and life history traits. Ecology 88:2662–2672

Garcia K, Lasco R, Ines A, Lyon B, Pulhin F (2013) Predicting geographic distribution and habitat suitability due to climate change of selected threatened forest tree species in the. Philippines. Appl Geogr 44:12–22

Halley JM, Monokrousos N, Mazaris AD, Newmark WD, Vokou D (2016) Dynamics of extinction debt across five taxonomic groups. Nat Commun 7:6

He XH, Duan YH, Chen YL, Xu MG (2010) A 60-year journey of mycorrhizal research in China: Past, present and future directions. Sci China-Life Sci 53:1374–1398

Hijmans RJ, Cameron SE, Parra JL, Jones PG, Jarvis A (2005) Very high resolution interpolated climate surfaces for global land areas. Int J Climatol 25:1965–1978

Huang JH, Ma KP, Huang JH (2017) Species Diversity Distribution Patterns of Chinese Endemic Seed Plants Based on Geographical Regions. PLoS One 12:13

Huberty CJ (1994) Applied Discriminant Analysis. Wiley Interscience, New York, USA.

Korner C (2007) The use of ‘altitude’ in ecological research. Trends Ecol Evol 22:569–574

Kraft NJB, Metz MR, Condit RS, Chave J (2010) The relationship between wood density and mortality in a global tropical forest data set. New Phytol 188:1124–1136

Kumar D, Rawat S (2022) Modeling the effect of climate change on the distribution of threatened medicinal orchid Satyrium nepalense D. Don in India. Environ Sci Pollut Res 29:72431–72444

Kuussaari M, Bommarco R, Heikkinen RK, Helm A, Krauss J, Lindborg R, Ockinger E, Partel M, Pino J, Roda F, Stefanescu C, Teder T, Zobel M, Steffan-Dewenter I (2009) Extinction debt: a challenge for biodiversity conservation. Trends Ecol Evol 24:564–571

Leigh A (2022) Using leaf shape to determine leaf size could be a game-changer. A commentary on: ‘Leaf size estimation based on leaf length, width and shape’. Ann Bot 129:I-II

Lenoir J, Gegout JC, Marquet PA, de Ruffray P, Brisse H (2008) A significant upward shift in plant species optimum elevation during the 20th century. Science 320:1768–1771

Li YQ, Reich PB, Schmid B, Shrestha N, Feng X, Lyu T, Maitner BS, Xu XT, Li YC, Zou DT, Tan ZH, Su XY, Tang ZY, Guo QH, Feng XJ, Enquist BJ, Wang ZH (2020) Leaf size of woody dicots predicts ecosystem primary productivity. Ecol Lett 23:1003–1013

Liao ZY, Zhang L, Nobis MP, Wu XG, Pan KW, Wang KQ, Dakhil MA, Du MX, Xiong QL, Pandey B, Tian XL (2020) Climate change jointly with migration ability affect future range shifts of dominant fir species in Southwest China. Divers Distrib 26:352–367

Lin LL, He J, Xie L, Cui GF (2020) Prediction of the Suitable Area of the Chinese White Pines (Pinus subsect. Strobus) under Climate Changes and Implications for Their Conservation. Forests 11:23

Liu M (1998) The atlas of the physical geography of China, 2nd end. Beijing: Sinomaps Press.

Lobell DB, Schlenker W, Costa-Roberts J (2011) Climate Trends and Global Crop Production Since 1980. Science 333:616–620

Lomba A, Pellissier L, Randin C, Vicente J, Moreira F, Honrado J, Guisan A (2010) Overcoming the rare species modelling paradox: A novel hierarchical framework applied to an Iberian endemic plant. Biol Conserv 143:2647–2657

Lu SF, Zhou SY, Yin XJ, Zhang C, Li RL, Chen JH, Ma DX, Wang Y, Yu ZX, Chen YH (2021) Patterns of tree species richness in Southwest China. Environ Monit Assess 193:13

Malcolm JR, Liu CR, Neilson RP, Hansen L, Hannah L (2006) Global warming and extinctions of endemic species from biodiversity hotspots. Conserv Biol 20:538–548.

Mao L, Swenson NG, Sui X, Zhang J, Chen S, Li J, Peng P, Zhou G, Zhang X (2019) The geographic and climatic distribution of plant height diversity for 19,000 angiosperms in China. Biodiversity and Conservation 29: 487–502.

Matern A, Drees C, Kleinwachter M, Assmann T (2007) Habitat modelling for the conservation of the rare ground beetle species Carabus variolosus (Cololeoptera, Carabidae) in the riparian zones of headwaters. Biol Conserv 136:618–627

Matteodo M, Wipf S, Stockli V, Rixen C, Vittoz P (2013) Elevation gradient of successful plant traits for colonizing alpine summits under climate change. Environ Res Lett 8:10

McGill BJ, Enquist BJ, Weiher E, Westoby M (2006) Rebuilding community ecology from functional traits. Trends Ecol Evol 21:178–185

Merow C, Smith MJ, Silander JA (2013) A practical guide to MaxEnt for modeling species’ distributions: what it does, and why inputs and settings matter. Ecography 36:1058–1069

Miller G, Magee J, Smith M, Spooner N, Baynes A, Lehman S, Fogel M, Johnston H, Williams D, Clark P, Florian C, Holst R, DeVogel S (2016) Human predation contributed to the extinction of the Australian megafaunal bird Genyornis newtoni similar to 47 ka. Nat Commun 7:7

Mueller TL, Karlsen-Ayala E, Moeller DA, Bellemare J (2022) Of mutualism and migration: will interactions with novel ericoid mycorrhizal communities help or hinder northward Rhododendron range shifts? Oecologia 198:839–852

Muscarella R, Galante PJ, Soley-Guardia M, Boria RA, Kass JM, Uriarte M, Anderson RP (2014) ENMeval: An R package for conducting spatially independent evaluations and estimating optimal model complexity for MAXENT ecological niche models. Methods Ecol Evol 5:1198–1205

Myers N, Mittermeier RA, Mittermeier CG, da Fonseca GAB, Kent J (2000) Biodiversity hotspots for conservation priorities. Nature 403:853-858

Nadeau CP, Fuller AK (2016) Combining landscape variables and species traits can improve the utility of climate change vulnerability assessments. Biol Conserv 202:30–38

Noh JK, Echeverria C, Pauchard A, Cuenca P (2019) Extinction debt in a biodiversity hotspot: the case of the Chilean Winter Rainfall-Valdivian Forests. Landsc Ecol Eng 15:1–12

O’Rourke H, Lughadha EN, Bacon KL (2022) Can the extinction risk of Irish vascular plants be predicted using leaf traits? Biodivers Conserv 31:3113–3135

Phillips OL, Sullivan MJP, Baker TR, Mendoza AM, Vargas PN, Vasquez R (2019) Species Matter: Wood Density Influences Tropical Forest Biomass at Multiple Scales. Surv Geophys 40:913–935

Phillips SJ, Anderson RP, Dudik M, Schapire RE, Blair ME (2017) Opening the black box: an open-source release of Maxent. Ecography 40:887–893

R Core Team (2022) R: A language and environment for statistical computing. R Foundation for Statistical Computing. Vienna, Austria. https://www.R-project.org

Record S, Charney ND, Zakaria RM, Ellison AM (2013) Projecting global mangrove species and community distributions under climate change. Ecosphere 4:23

Reich PB, Wright IJ, Cavender-Bares J, Craine JM, Oleksyn J, Westoby M, Walters MB (2003) The evolution of plant functional variation: Traits, spectra, and strategies. Int J Plant Sci 164:S143–S164

Roberts DL, Solow AR (2003) Flightless birds - When did the dodo become extinct? Nature 426:245–245

Saar L, Takkis K, Partel M, Helm A (2012) Which plant traits predict species loss in calcareous grasslands with extinction debt? Divers Distrib 18:808–817

Schrader J, Shi PJ, Royer DL, Peppe DJ, Gallagher RV, Li YR, Wang R, Wright IJ (2021) Leaf size estimation based on leaf length, width and shape. Ann Bot 128:395–406

Seto KC, Kaufmann RK, Woodcock CE (2000) Landsat reveals China’s farmland reserves, but they’re vanishing fast. Nature 406:121–121

Shi XD, Yin Q, Sang ZY, Zhu ZL, Jia ZK, Ma LY (2021) Prediction of potentially suitable areas for the introduction of Magnolia wufengensis under climate change. Ecol Indic 127:14

Skov F, Svenning JC (2004) Potential impact of climatic change on the distribution of forest herbs in Europe. Ecography 27:366–380

Stuecker MF, Timmermann A, Jin FF, Proistosescu C, Kang SM, Kim D, Yun KS, Chung ES, Chu JE, Bitz CM, Armour KC, Hayashi M (2020) Strong remote control of future equatorial warming by off-equatorial forcing. Nat Clim Chang 10:124

Swets JA (1988) Measuring the accuracy of diagnostic systems. Science 240:1285–1293

Thomas CD, Cameron A, Green RE, Bakkenes M, Beaumont LJ, Collingham YC, Erasmus BFN, de Siqueira MF, Grainger A, Hannah L, Hughes L, Huntley B, van Jaarsveld AS, Midgley GF, Miles L, Ortega-Huerta MA, Peterson AT, Phillips OL, Williams SE (2004) Extinction risk from climate change. Nature 427:145–148

Thomson FJ, Letten AD, Tamme R, Edwards W, Moles AT (2018) Can dispersal investment explain why tall plant species achieve longer dispersal distances than short plant species? New Phytol 217:407–415

Thurm EA, Hernandez L, Baltensweiler A, Ayan S, Rasztovits E, Bielak K, Zlatanov TM, Hladnik D, Balic B, Freudenschuss A, Buchsenmeister R, Falk W (2018) Alternative tree species under climate warming in managed European forests. For Ecol Manage 430:485–497

Tilman D, May RM, Lehman CL, Nowak MA (1994) Habitat destruction and the extinction debt. Nature 371:65–66

Tomiolo S, Ward D (2018) Species migrations and range shifts: A synthesis of causes and consequences. Perspectives in Plant Ecology, Evolution and Systematics 33: 62–77.

Trenberth KE (2011) Changes in precipitation with climate change. Clim Res 47:123–138

Vellend M, Verheyen K, Jacquemyn H, Kolb A, Van Calster H, Peterken G, Hermy M (2006) Extinction debt of forest plants persists for more than a century following habitat fragmentation. Ecology 87:542–548

Veresoglou SD, Halley JM (2018) Seed mass predicts migration lag of European trees. Ann For Sci 75:9

Wang Y, Ma XH, Lu YF, Hu XE, Lou LH, Tong ZK, Zhang JH (2022) Assessing the current genetic structure of 21 remnant populations and predicting the impacts of climate change on the geographic distribution of Phoebe sheareri in southern China. Sci Total Environ 846:13

Warren DL, Matzke NJ, Cardillo M, Baumgartner JB, Beaumont LJ, Turelli M, Glor RE, Huron NA, Simoes M, Iglesias TL, Piquet JC, Dinnage R (2021) ENMTools 1.0: an R package for comparative ecological biogeography. Ecography 44:504–511

Warren DL, Seifert SN (2011) Ecological niche modeling in Maxent: the importance of model complexity and the performance of model selection criteria. Ecol Appl 21:335–342

Wright IJ, Falster DS, Pickup M, Westoby M (2006a) Cross-species patterns in the coordination between leaf and stem traits, and their implications for plant hydraulics. Physiol Plant 127:445–456

Wright IJ, Leishman MR, Read C, Westoby M (2006b) Gradients of light availability and leaf traits with leaf age and canopy position in 28 Australian shrubs and trees. Funct Plant Biol 33:407–419

Yang JT, Jiang P, Huang Y, Yang YL, Wang RL, Yang YX (2022) Potential geographic distribution of relict plant Pteroceltis tatarinowii in China under climate change scenarios. PLoS One 17:21

Yang XQ, Kushwaha SPS, Saran S, Xu JC, Roy PS (2013) Maxent modeling for predicting the potential distribution of medicinal plant, Justicia adhatoda L. in Lesser Himalayan foothills. Ecol Eng 51:83–87

Yigini Y, Panagos P (2016) Assessment of soil organic carbon stocks under future climate and land cover changes in Europe. Sci Total Environ 557:838–850

Zhang KL, Yao LJ, Meng JS, Tao J (2018) Maxent modeling for predicting the potential geographical distribution of two peony species under climate change. Sci Total Environ 634:1326–1334

Zhao YT, Ali A, Yan ER (2017) The plant economics spectrum is structured by leaf habits and growth forms across subtropical species. Tree Physiol 37:173–185

## References

Dormann, C. F., Elith, J., Bacher, S., Buchmann, C., Carl, G., Carre, G., . . . Lautenbach, S. (2013). Collinearity: a review of methods to deal with it and a simulation study evaluating their performance. Ecography, 36(1), 27–46. doi:10.1111/j.1600-0587.2012.07348.x

Garcia, K., Lasco, R., Ines, A., Lyon, B., & Pulhin, F. (2013). Predicting geographic distribution and habitat suitability due to climate change of selected threatened forest tree species in the. Philippines. Applied Geography, 44, 12–22. doi:10.1016/j.apgeog.2013.07.005

Merow, C., Smith, M. J., & Silander, J. A. (2013). A practical guide to MaxEnt for modeling species’ distributions: what it does, and why inputs and settings matter. Ecography, 36(10), 1058–1069. doi:10.1111/j.1600-0587.2013.07872.x

Naimi, B., Hamm, N. A. S., Groen, T. A., Skidmore, A. K., & Toxopeus, A. G. (2014). Where is positional uncertainty a problem for species distribution modelling? Ecography, 37(2), 191–203. doi:10.1111/j.1600-0587.2013.00205.x

Phillips, S. J. (2008). Transferability, sample selection bias and background data in presence-only modelling: a response to Peterson et al. (2007). Ecography, 31(2), 272-278. doi:10.1111/j.0906-7590.2008.5378.x

